# Geranylgeranyl Isoprenoids and Hepatic Rap1a Regulate Basal and Statin-Induced Expression of PCSK9

**DOI:** 10.1101/2023.10.23.563509

**Authors:** Yating Wang, Brea Tinsley, Stefano Spolitu, John A Zadroga, Heena Agarwal, Amesh K Sarecha, Lale Ozcan

## Abstract

Low-density lipoprotein cholesterol (LDL-C) lowering is the main goal of atherosclerotic cardiovascular disease prevention, and proprotein convertase subtilisin/kexin type 9 (PCSK9) inhibition is now a validated therapeutic strategy that lowers serum LDL-C and reduces coronary events. Ironically, the most widely used medicine to lower cholesterol, statins, has been shown to increase circulating PCSK9 levels, which limits their efficacy. Here, we show that geranylgeranyl isoprenoids and hepatic Rap1a regulate both basal and statin induced expression of PCSK9 and contribute to LDL-C homeostasis. Rap1a prenylation and activity is inhibited upon statin treatment, and statin mediated PCSK9 induction is dependent on geranylgeranyl synthesis and hepatic Rap1a. Accordingly, treatment of mice with a small molecule activator of Rap1a lowered PCSK9 protein and plasma cholesterol and inhibited statin mediated PCSK9 induction in hepatocytes. The mechanism involves inhibition of the downstream RhoA-ROCK pathway and regulation of PCSK9 at the post transcriptional level. These data further identify Rap1a as a novel regulator of PCSK9 protein and show that blocking Rap1a prenylation through lowering geranylgeranyl levels contributes to statin-mediated induction of PCSK9.

## Introduction

Lipid lowering therapies remain the cornerstone treatment of atherosclerosis and continual improvements in therapies are still at the forefront of discovery. PCSK9 inhibitors emerged as a novel treatment approach to lower serum LDL-C and are well-tolerated.^1,2^ PCSK9 binds to LDL receptor (LDLR) and increases its lysosomal degradation, which results in decreased rate of LDL-C removal from the circulation.^3^ PCSK9 may also contribute to atherosclerosis development through its effects on endothelial and vascular smooth muscle cells.^4,5^ Accumulating evidence suggest that PCSK9 inhibition has many beneficial effects besides LDL-C management, including lowering of atherogenic Lp(a) levels and protecting against calcific aortic valve stenosis.^6–8^ Anti-PCSK9 antibodies lower LDL-C and reduce coronary events; however, their high costs and low patient compliance limit their use.^9^ Liver-targeted siRNA against PCSK9 also robustly lowers LDL-C, but unlike the well-tolerated genetic variants, it removes all PCSK9 from the cell, raising safety concerns.^10^ Thus, further studies aimed at identifying the mechanism(s) of PCSK9 regulation are crucial, which could be used to develop new therapeutic strategies.

Statins inhibit HMG-CoA reductase, which catalyzes the rate-limiting step in cholesterol synthesis, and upregulate LDLR.^11^ Although statins are the most widely-prescribed drugs to lower cholesterol, a substantial proportion of patients do not reach target LDL-C goals despite receiving maximally tolerated statin therapy.^12–15^ In fact, each doubling of the statin dose was shown to produce an average additional decrease in LDL-C levels of about only 6% (rule of 6)^16,17^. This was attributed in part to an increase in PCSK9 levels upon statin use, which promotes LDLR degradation and decreases statins’ LDL-C lowering efficacy.^18–22^ Accordingly, combination of anti-PCSK9 antibodies with statins provides 50-60% additional reduction in LDL-C and reduces cardiovascular events by 50%.^23^ On the other hand, clinical trials of selective cholesterol absorption inhibitor, ezetimibe, have not shown an effect on plasma PCSK9 levels, suggesting that PCSK9 induction by statins may not be simply attributed to their effects on plasma LDL-C.^20,24^ Thus, there is a need to understand the underlying molecular mechanisms of this ironic relationship between statins and plasma PCSK9 that can lay a path to improve their efficacy. An increase in nuclear localization of sterol regulatory element (SRE)-binding protein 2 (SREBP2), which binds to the SRE motifs located in the promoter region *of Pcsk9* gene, has been shown as one mechanism by which statins adversely increase PCSK9 levels.^25,26^ It is unknown, however, whether there are additional pathways involved in statin-mediated PCSK9 induction. In addition to lowering cholesterol synthesis and regulating LDL-C levels, statins also inhibit the formation of other mevalonate pathway metabolites, including farnesyl pyrophosphate (FPP) and geranylgeranyl pyrophosphate (GGPP) isoprenoids, which are necessary for the prenylation and activation of small G proteins, including Rap1a.^27–29^

Rap1a is a small GTPase, which gets activated by GTP loading.^30–32^ Attachment of GGPP isoprenoid chain to Rap1a by geranylgeranyl transferase-I (GGTase-I) results in Rap1a’s membrane localization and its activation by guanine exchange factors, including exchange factor activated by cAMP-2 (Epac2).^33,34^ The GTPase-activating protein, Rap1GAP, hydrolyzes Rap1a’s GTP and inactivates it. Our previous work revealed that activation of glucagon receptor–cAMP–Rap1a pathway in the liver decreases hepatic and plasma PCSK9, which leads to increased LDLR protein and lower plasma LDL-C.^35^ Specifically, we reported that silencing of Rap1a in isolated hepatocytes increases PCSK9 and acute inhibition of Rap1a activity in mice liver via adenovirus mediated-overexpression of Rap1GAP elevates plasma PCSK9, lowers hepatic LDLR and increases LDL-C. Our mechanistic studies revealed that Rap1a decreases the stability of PCSK9 protein without changing its mRNA or SREBP2 activity.^35^ In addition to its LDL-C lowering effect, we recently showed that activation of Rap1a suppresses hepatic glucose production and its inhibition contributes to obesity-induced glucose intolerance.^36^ Further, statins decrease Rap1a prenylation and activity in isolated hepatocytes and human liver samples and this may in turn explain their hyperglycemic effects.^36^

Because Rap1a activity is decreased upon statins and adding back geranylgeranyl isoprenoids restores hepatic Rap1a activity, we became interested in the role of GGPP and Rap1a in statin-mediated PCSK9 induction.^36^ In this study, we show that restoration of GGPP levels in statin treated mice lowers statin-induced PCSK9 and further decreases plasma LDL-C, independent of a change in *Pcsk9* mRNA. Moreover, treatment of mice with a small molecule activator of Rap1a recapitulates the Rap1a gain-of-function model and lowers plasma PCSK9 and cholesterol levels. Rap1a activator treatment also lowers statin induced PCSK9 in hepatocytes. We further report that Rap1a regulates PCSK9 via inhibiting the RhoA-ROCK pathway. These findings provide an additional mechanistic insight into the paradoxical regulation of PCSK9 by statins and reveal that GGPP─Rap1a pathway contributes to the post-transcriptional regulation of PCSK9.

## Materials and Methods

### Mouse Experiments

WT (stock # 000664) and diet-induced obese (DIO, stock # 380050) mice were from Jackson Labs. DIO mice were fed a high-fat diet with 60% kcal from fat (Research Diets). To avoid the effects of gender-related differences in metabolism, male mice were used for most of the experiments. All mice were maintained on a 12 h-light-dark cycle. Recombinant adeno-associated viruses (AAVs, 1-2 X 10^9^ genome copies per mouse) and adenoviruses (0.75 X 10^9^ plaque-forming units per mouse) were delivered to DIO mice by tail vein injections, and experiments were commenced after 3-28 days. In some experiments, high cholesterol containing diet fed WT mice were used, as indicated in the corresponding legends.^37^ For some of the statin treatment experiments, 18-week-old male DIO mice were treated orally with rosuvastatin (10 mg/kg/day) for 3 weeks or 0.02% (w/w) simvastatin was mixed with the high-fat diet. Geranylgeraniol (GGOH) was given by oral gavage at a dose of 50 mg/kg/day for 3 weeks. 20-week-old male DIO mice were treated intraperitoneally with 8-pCPT 1.5 mg/kg/day for 2 weeks. For all experiments, male mice of the same age and similar weight were randomly assigned to experimental and control groups. Animal studies were conducted in accordance with the Columbia University Irving Medical Center Animal Care and Use Committee.

### Reagents and Antibodies

Rosuvastatin (cat # 18813), simvastatin for cell culture (cat # 10010344), GGPP (cat # 63330) and GGOH (cat # 13272) were from Cayman Chemicals. C3 (Rho inhibitor I, cat # CT04) was from Cytoskeleton. Anti-β-actin (cat # 4970), anti-Hsp90 (cat # 4877), anti-RhoA (cat # 21117), anti-Gapdh (cat # 5174), and anti-pan-Cadherin (cat # 4068) antibodies were from Cell Signaling Technology. Anti-PCSK9 antibody was from R&D Systems (catalog # AF3985). Specific Epac activator, 8-pCPT, was from Cayman Chemicals (cat # 17143). AAV8-shRNA targeting murine *Pggt1b* was made by annealing complementary oligonucleotides 5’-CACCAAAGCCATCAGCTACATTAGAAGAAGTCAAGAGCTTCTTCTAATGTAGCTGATGGCTT −3’, which were then ligated into the self-complementary AAV8-RSV-GFP-H1 vector as described previously.^38^ The resultant constructs were amplified by Salk Institute Gene Transfer, Targeting, and Therapeutics Core. AAV8-sh-RhoA was as described before.^37^ siRNA sequences against murine *Rap1a*, *Pggt1b*, *Rhoa* and *Rock* were purchased from Integrated DNA Technologies. Adenoviruses encoding LacZ and Rap1Gap was described previously^31,39^ and amplified and purified by Viraquest, Inc. (North Liberty, IA). Myc-tagged WT Rap1a or my-tagged prenylation deficient Rap1a (SAAX-Rap1a) plasmids were gifts from Dr. Carol L. Williams (Medical College of Wisconsin).^40^

### Measurement of lipids, insulin, and glucose

Plasma total cholesterol levels were measured using a colorimetric kit (Wako). Plasma lipoproteins were separated by KBr density ultracentrifugation as described previously.^41^ Briefly, equal amounts of mouse plasma (50–100 μL) were used for sequential density ultracentrifugation to separate very-low-density lipoprotein (d<1.006 g/mL), low-density lipoprotein (LDL, d=1.006–1.063 g/mL), and high-density lipoprotein (d=1.063–1.21 g/mL) in a TLA 100 rotor and LDL fractions were used to measure cholesterol levels using the above colorimetric kit. Plasma PCSK9 levels were analyzed using an ELISA kit (R&D Systems).

### Hepatocyte experiments

Primary mouse hepatocytes were isolated from 12-20-week-old male or female WT mice as described previously.^42^ In some experiments, primary mouse hepatocytes were isolated from hepatocyte-specific Rap1a deficient mice that were obtained by injecting *Rap1a*^fl/fl^ mice (Jackson Labs, stock number: 021066) with AAV expressing Cre recombinase, driven by the thyroxin-binding globulin (TBG) promoter (AAV8-TBG-Cre). Hepatocytes were transfected with plasmids 10-12 hr after plating, and experiments were conducted 48-72 hr later. Transfections with scrambled, si-Rap1a, si-Pggt1b, si-RhoA or si-ROCK constructs were carried out using Lipofectamine RNAiMAX reagent according to manufacturer’s instructions.

### Immunoblotting

Liver protein was extracted using RIPA buffer (Thermo Scientific), and the protein concentration was measured by DC protein assay kit (Bio-Rad). Whole cell lysates were harvested in 2X Laemmli buffer. Cell extracts were electrophoresed on SDS-polyacrylamide gels and transferred to 0.2 μm or 0.45 μm PVDF membranes. Blots were blocked in Tris-buffered saline with 0.1% Tween-20 containing 5% BSA at room temperature for one hour. Membranes were then incubated overnight at 4°C with primary antibodies. The protein bands were detected with horseradish peroxidase-conjugated secondary antibodies (Jackson ImmunoResearch) and Supersignal West Pico enhanced chemiluminescent solution (Thermo).

### Rap1a Activity Assay

Rap1a activity was assayed using Active Rap1 Detection Kit (Cell Signaling Technology, cat # 8818). Liver tissue sample were lysed with lysis/binding/washing buffer containing 2 mM PMSF, 5 µg/ml leupeptin, 10 nM okadaic acid and 5 µg/ml aprotinin. GST-RalGDS-RBD fusion protein was used to bind the activated form of GTP-bound Rap1a, which was then immunoprecipitated with glutathione resin. Rap1a activation levels were determined by western blot using a Rap1a primary antibody (R&D Systems, cat # AF3767).

### Quantitative PCR

RNA was isolated from hepatocytes or liver tissue using TRIzol (Invitrogen) and cDNA was synthesized from 1 ug of RNA using a cDNA synthesis kit (Invitrogen). Real-time PCR was performed using a 7500 Real-Time PCR system and SYBR Green reagents (Applied Biosystems). Specific primer sets used were as follows*: Pcsk9* forward: GAGACCCAGAGGCTACAGATT, *Pcsk9* reverse: AATGTACTCCACATGGGGCAA; *Rplp0* (36B4-housekeeping) forward: GCTCCAAGCAGATGCAGCA, *Rplp0* (36B4-housekeeping) reverse: CCGGATGTGAGGCAGCAG

### Statistical analysis

All results are presented as mean ± SEM. *P* values were calculated using the Student’s *t*-test for normally distributed data and the Mann-Whitney rank sum test for non-normally distributed data. For experiments with more than two groups, *P* values were calculated using one-way ANOVA for normally distributed data and the Kruskal-Wallis by ranks test for non-normally distributed data.

## Results

### Geranylgeranyl isoprenoid restoration suppresses statin induced PCSK9 independent of a change in *Pcsk9* mRNA and lowers LDL-C

Inhibition of HMG-CoA reductase by statins leads to reduced production of FPP and GGPP isoprenoids. Because GGPP is required for Rap1a activation, and Rap1a is an important regulator of PCSK9,^35^ we reasoned that a decrease in intracellular GGPP levels might be involved in statin mediated PCSK9 induction. To test this, we restored intracellular GGPP in statin-treated hepatocytes by incubating them with exogenous GGPP, which we showed previously to bring back Rap1a’s prenylation and plasma membrane localization to control cells.^36^ Consistent with restoration of Rap1a’s membrane localization, GGPP treatment significantly lowered PCSK9 protein in statin-treated hepatocytes (**Figure 1A**). Importantly, GGPP treatment did not inhibit simvastatin-induced mRNA expression of *Pcsk9* (**Figure 1B**) or other SREBP2 target genes, suggesting that activation of SREBP2 pathway upon statin treatment was not altered by GGPP restoration.^36^ We next investigated the functional importance in vivo and treated diet-induced obese WT mice with 10 mg/kg/day rosuvastatin alone or in combination with 50 mg/kg/day geranylgeraniol (GGOH) by daily gavage for 3 weeks. GGOH is an intermediate product in the mevalonate pathway and acts as a precursor to GGPP.^43^ Upon GGOH add back, hepatic Rap1a activity was restored and statin-induced hepatic and plasma PCSK9 levels were lowered (**Figure 1C-E**). As WT mice are resistant to statins’ LDL-C lowering effect due in part to strong induction of PCSK9 by statins,^44^ rosuvastatin alone had no effect on plasma LDL-C (**Figure 1F**). However, when PCSK9 levels were lowered upon GGPP restoration, plasma LDL-C was decreased in statin-treated mice (**Figure 1F**), without a change in body weight or food intake (not shown). These results support the hypothesis that inhibition of the mevalonate pathway by statins increase circulating PCSK9 in part via lowering intracellular GGPP levels, which is independent of the SREBP2-mediated *Pcsk9* mRNA induction.

**Figure 1.**
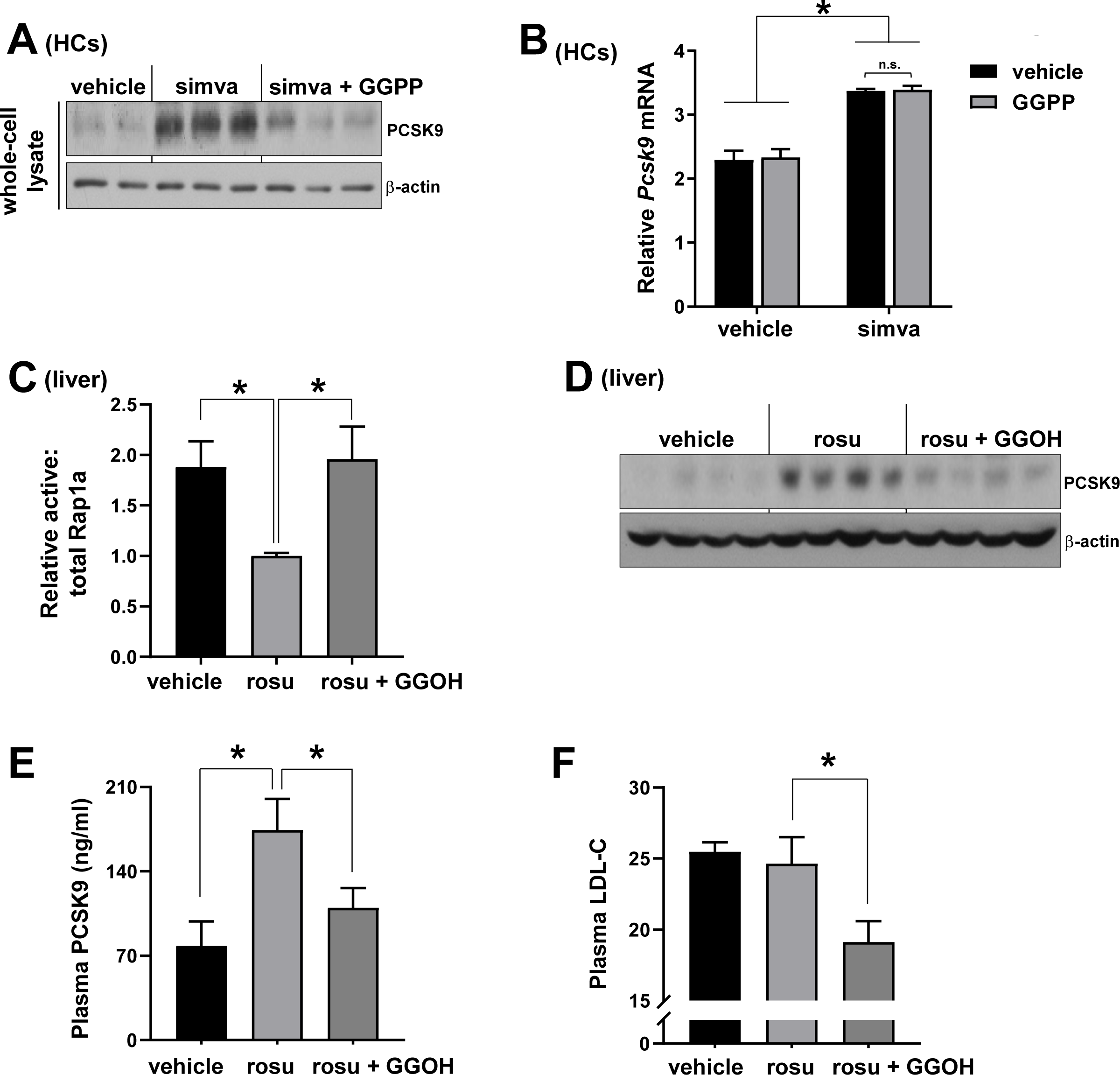
Geranylgeranyl isoprenoid restoration suppresses statin induced PCSK9 independent of a change in *Pcsk9* mRNA and lowers LDL-C. **(A-B)** Primary hepatocytes (HCs) were treated with vehicle, 10 μM simvastatin (simva) or simvastatin and 10 μM geranylgeranyl pyrophosphate (GGPP) together for 32 hours. PCSK9 protein (A) and *Pcsk9* mRNA (B) levels were measured (n=2-4, mean ± SEM, \**P* <0.05, n.s., non-significant). **(C-F)** Densitometric quantification of hepatic active:total Rap1a (C), liver PCSK9 protein (D), plasma PCSK9 (E) and plasma LDL-C (F) levels in high-fat diet fed WT C57BL/6J male mice that were treated with vehicle, 10 mg/kg rosuvastatin (rosu) or rosu + geranylgeraniol (GGOH) (50 mg/kg) daily for 3 weeks (n=4-5, mean ± SEM, *p < 0.05).

### GGTase-I inhibition and prenylation deficient Rap1a mimic statins and induce PCSK9 protein under basal conditions

We next silenced GGTase-I, which catalyzes the covalent attachment of GGPP to the cysteine residue in the C-terminal CAAX motif of Rap1a to form a stable thioether bond.^45–47^ Similar to statin treatment or Rap1a inhibition, silencing *Pggt1b*, the gene encoding the GGTase-I enzyme, in hepatocytes increased both intracellular and secreted PCSK9 without affecting *Pcsk9* mRNA (**Figure 2A-2B**).In agreement with the in vitro data, silencing Pggt1b in the liver by treatment of WT mice with the hepatocyte-specific AAV8-H1-shPggt1b increased plasma PCSK9 and had no effect on hepatic *Pcsk9* mRNA levels (**Figure 2C-2D**). We then asked whether the prenylation defective mutant form of Rap1a mimics statin treatment or GGTase-I deficiency and induces PCSK9. We transfected Rap1a KO hepatocytes with an empty plasmid or equal amounts of plasmids encoding the WT-or prenylation-deficient mutant form of Rap1a where the cysteine in the CAAX motif is replaced with serine to generate the non-prenylated Rap1a-SAAX mutant (**Figure 2E**).^40^ As expected, reconstituting WT Rap1a lowered PCSK9 protein as compared to empty plasmid treated Rap1a KO cells, whereas Rap1a-SAAX mutant treated cells expressed higher levels of PCSK9 protein without a change in *Pcsk9* mRNA (**Figure 2F-2G**). These data suggest the idea that inhibition of Rap1a prenylation induces PCSK9, which could contribute to the statin-PCSK9 link.

**Figure 2.**
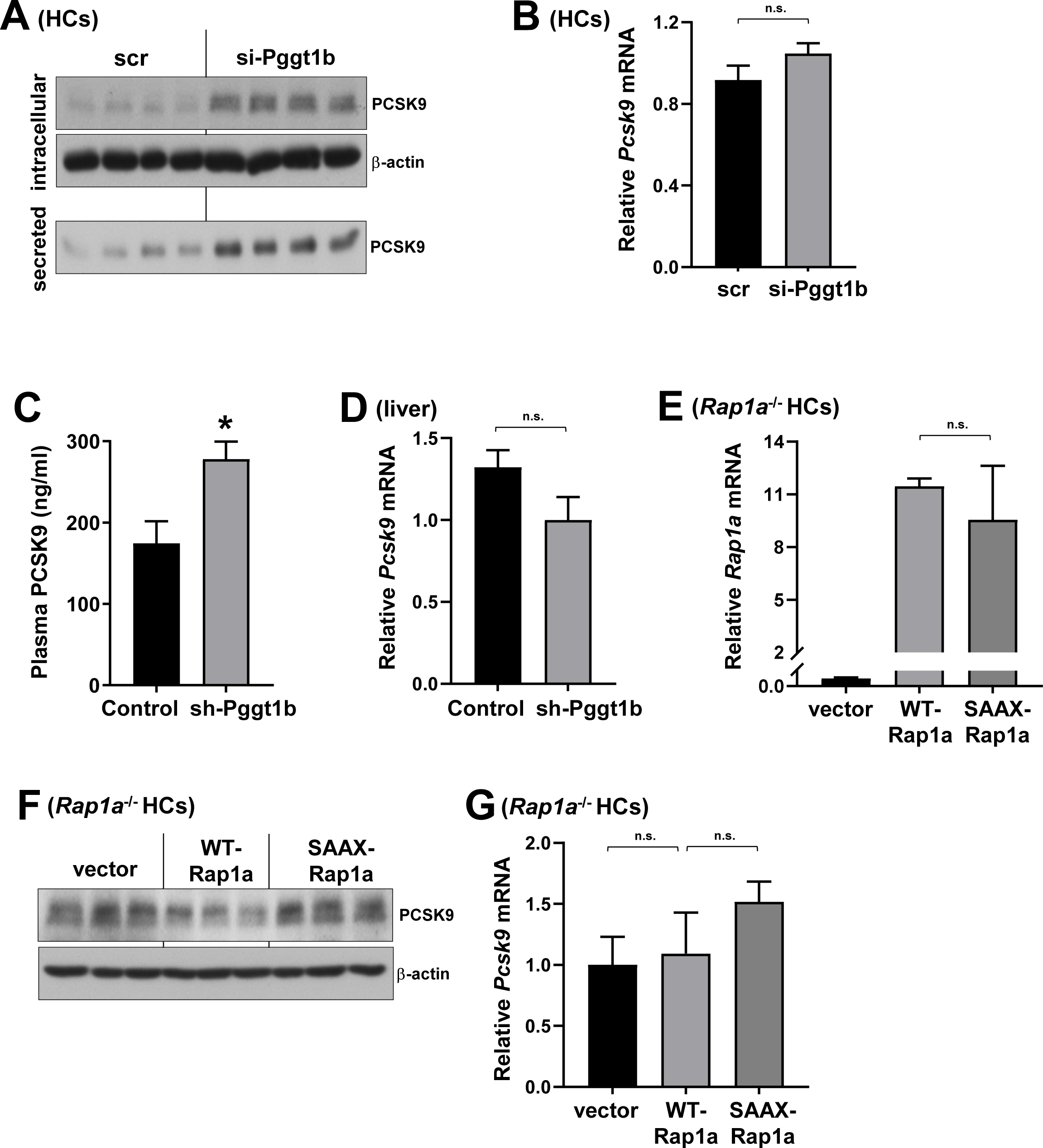
GGTase-I silencing and prenylation deficient-Rap1a increase PCSK9. (**A-B**) Intracellular and secreted PCSK9 protein (A) or *Pcsk9* mRNA (B) from scrambled (scr) or si-Pggt1b–treated primary hepatocytes (HCs). **(C-D)** Plasma PCSK9 (C) and liver *Pcsk9* mRNA levels (D) from high-fat diet fed WT C57BL/6J male mice treated with an AAV8-H1 construct encoding sh-Pggt1b or empty vector (control) (n=7-8/group, mean ± SEM, \**P* <0.05). **(E-G)** *Rap1a*^-/-^hepatocytes were transfected with empty plasmid (vector) or plasmids encoding WT or prenylation-deficient Rap1a (SAAX-Rap1a), and *Rap1a* mRNA (E), intracellular PCSK9 (F) and *Pcsk9* mRNA levels were assayed (G).

### Rap1a activation via 8-pCPT treatment suppresses basal and statin-induced PCSK9

Because our previous work showed that overexpression of a constitutively active Rap1a lowers plasma PCSK9,^35^ we asked whether activating Rap1a pharmacologically could have similar beneficial effects. The Epac activator, 8-pCPT-2’-O-Me-cAMP (8-pCPT), is a cAMP analog, where the 2′-*O*-methyl and 8-chlorophenylthio groups confer specific Epac binding and robustly activate Rap1a and our previous work revealed that 8-pCPT treatment lowers PCSK9 secretion in isolated hepatocytes.^35,48^ We next tested whether 8-pCPT can lower circulating PCSK9 in vivo. Based on the literature and our in vitro potency data, we treated high-fat diet fed mice with 1.5 mg/kg body wt i.p. 8-pCPT daily for two weeks. The mice showed no signs of toxicity, and there were no differences in body weight or food intake by 8-pCPT (not shown). Rap1a activity in the liver was increased as evidenced by higher GTP-bound Rap1a levels (**Figure 3A**), and as predicted by our hypothesis, 8-pCPT treatment lowered both plasma PCSK9 and total cholesterol levels (**Figure 3B-3C**). We then asked whether forced Rap1a activation via 8-pCPT treatment can overcome the statin mediated Rap1a inhibition and lower PCSK9. We found that 8-pCPT pretreatment of isolated hepatocytes lowered simvastatin induced intracellular and secreted PCSK9 protein (**Figure 3D**). Collectively, these data provide additional support for the hypothesis that statins increase PCSK9 in part by inhibiting the GGPP–Rap1a pathway in hepatocytes.

**Figure 3.**
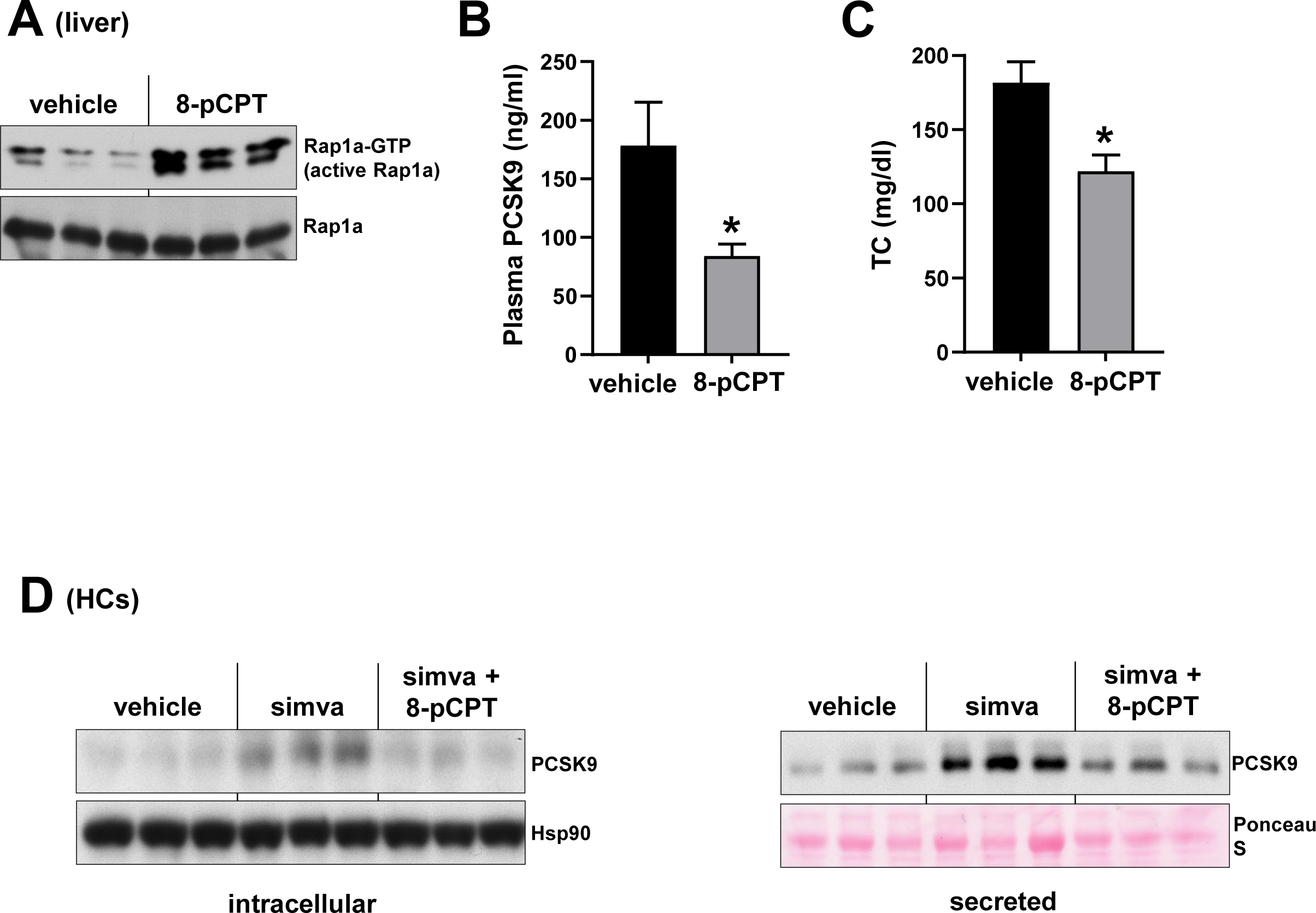
Rap1a activation via 8-pCPT treatment suppresses basal and statin-induced PCSK9. **(A-C)** WT C57BL/6J male mice that were fed with high-fat diet were treated with 1.5 mg/kg/day with 8-pCPT for 2 weeks. Hepatic GTP-Rap1a (active Rap1a) (A), plasma PCSK9 (B), and plasma total cholesterol (TC) (C) were measured (n= 4 mice/group, mean ± SEM, *p < 0.05). **(D)** Intracellular (left panel) and secreted PCSK9 (right panel) levels were assayed in primary hepatocytes (HCs) treated with vehicle, 10 μm simvastatin (simva) or simvastatin and 20 μm 8-pCPT together for 48 hours.

### Rap1a silencing increases PCSK9 via inducing the RhoA-ROCK pathway

RhoA is a ubiquitously expressed small G protein, which gets activated by GTP loading.^49^ Although geranylgeranylation of RhoA is important for its plasma membrane localization, statin administration or inhibition of GGTase-I, which prenylates RhoA, paradoxically increases the protein expression and activity of RhoA in many cell types, including the hepatocytes.^50–54^ Similar to statin treatment, inhibition of Rap1a has been shown to induce RhoA and increase its active-GTP-bound form in endothelial cells.^55,56^ In light of the proposed role of RhoA in LDLR regulation,^54^ we hypothesized that RhoA inhibition could be a key downstream event by which Rap1a regulates PCSK9 and LDLR. Supporting this idea, we first showed that Rap1a silencing in isolated hepatocytes increased RhoA levels (**Figure 4A**). Correspondingly, we found that liver adeno-Rap1aGAP overexpression, which inhibits Rap1a activity, increased RhoA protein, suggesting that Rap1a → ↓RhoA axis exists in liver (**Figure 4B**). To test the importance of RhoA in PCSK9 and LDLR regulation, we silenced it in isolated hepatocytes. Similar to Rap1a activation, we observed that both intracellular and secreted PCSK9 were decreased upon si-RhoA treatment, without an effect on the *Pcsk9* mRNA (**Figure 4C-4D**). Additionally, RhoA inhibition using the RhoA-specific inhibitor, C3, also lowered PCSK9 in hepatocytes (**Figure 4E**). In order to assess the in vivo role of RhoA, we analyzed plasma and liver samples from control vs RhoA-silenced livers using hepatocyte-specific AAV8-H1-shRhoA^37^ and found that liver RhoA silencing decreased plasma PCSK9 and plasma cholesterol levels (**Figure 4F-4G**). Next, we asked whether the increase in PCSK9 upon Rap1a inhibition is dependent on RhoA. To test this, we silenced RhoA in si-Rap1a–treated hepatocytes to decrease RhoA protein to a level similar to that in control cells. The data showed that the effect of Rap1a silencing on PCSK9 induction was abrogated when RhoA was also silenced (**Figure 4H**), which suggest that the dominant pathway by which Rap1a inhibition induces PCSK9 is via increasing RhoA. We also tested whether the well-known RhoA effector, ROCK, participates in PCSK9 regulation and found that the ability of RhoA silencing to decrease intracellular and secreted PCSK9 was mimicked by silencing ROCK (**Figure 4I**).

**Figure 4.**
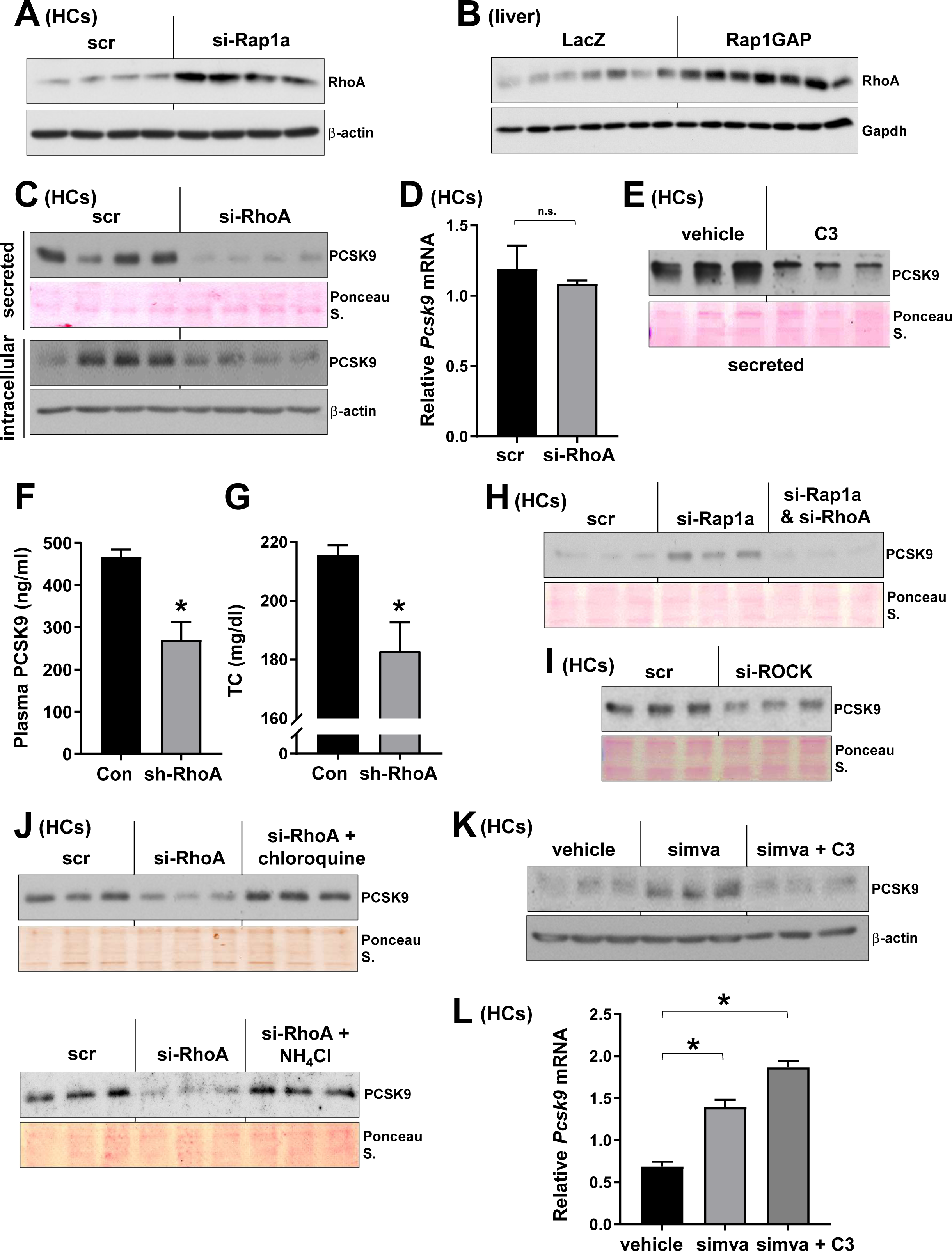
Rap1a silencing increases PCSK9 via inducing the RhoA-ROCK pathway. **(A-B)** RhoA levels were assayed in primary hepatocytes (HCs) treated with scrambled (scr) vs si-Rap1a (A), or in adeno-LacZ vs adeno-Rap1aGAP-expressing mice liver (B). **(C-D)** Scramble (scr) or si-RhoA–treated mouse hepatocytes were assayed for secreted and intracellular PCSK9 (C) or *Pcsk9* mRNA (D). (**E**) Mouse hepatocytes were treated with vehicle or 1 μg/mL C3 transferase Rho inhibitor (C3) and secreted PCSK9 levels were assayed. **(F-G)** Plasma PCSK9 (F) and plasma total cholesterol (TC) levels (G) were measured from AAV8-sh-RhoA or control AAV (Con)-treated mice fed with a high-cholesterol diet (n=3-4/group, mean ± SEM, \**P* <0.05). **(H**) Scramble (scr), si-Rap1a alone, or si-Rap1a + si-RhoA–treated primary hepatocytes were assayed for secreted PCSK9. (**I**) Same as in (A), except that scr- and si-ROCK– treated cells were used. **(J)** Secreted PCSK9 was assayed from scramble (scr) or si-RhoA–expressing cells that were treated with chloroquine (upper panel) or ammonium chloride (NH_4_Cl, lower panel). **(K-L)** Intracellular PCSK9 protein (K) and *Pcsk9* mRNA (L) levels from primary hepatocytes treated with vehicle, 10 mM simvastatin (simva) and simva + 1 μg/mL C3 for 32 hours.

Previous studies revealed that PCSK9 gets degraded in the lysosomes and our published work showed that activation of Rap1a increased the lysosomal degradation of PCSK9.^35,57,58^ Correspondingly, we observed that disruption of lysosomal function via chloroquine or ammonium chloride treatment prevented the decrease in PCSK9 protein conferred by RhoA silencing (**Figure 4J**), suggesting that RhoA inhibition, similar to Rap1a activation, enhances the lysosomal degradation of PCSK9. Of note, inhibition of RhoA-ROCK pathway did not affect hepatic albumin or alpha-1-antitrypsin secretion indicating that general protein secretion was not altered upon RhoA or ROCK silencing (not shown). Finally, we found that RhoA-silencing decreased statin-induced PCSK9 (**Figure 4K**) without an effect on *Pcsk9* mRNA (**Figure 4L**), which recapitulates GGPP restoration and Rap1a activator results. Altogether these data suggest the existence of Rap1a → ↓RhoA/ROCK → ↓PCSK9 axis in hepatocytes that could contribute to LDL-C regulation.

## Discussion

One mechanism by which statins paradoxically increase plasma PCSK9 levels is via inducing the SREBP2 pathway, which regulate PCSK9 at the mRNA level. Here we report an additional mechanism that involves inhibition of GGPP synthesis and Rap1a activity, which upregulates PCSK9 post transcriptionally via the RhoA-ROCK pathway (**Figure 5**). The data suggest that the same Rap1a-RhoA-ROCK pathway also regulates basal PCSK9 levels and can therapeutically be targeted via treatment with an Rap1a activator. In this regard previous work has shown that the small molecule activator of Rap1, 8-pCPT, protects against ischemia-reperfusion injury-induced renal failure and asthmatic airway inflammation.^59,60^

**Figure 5.**
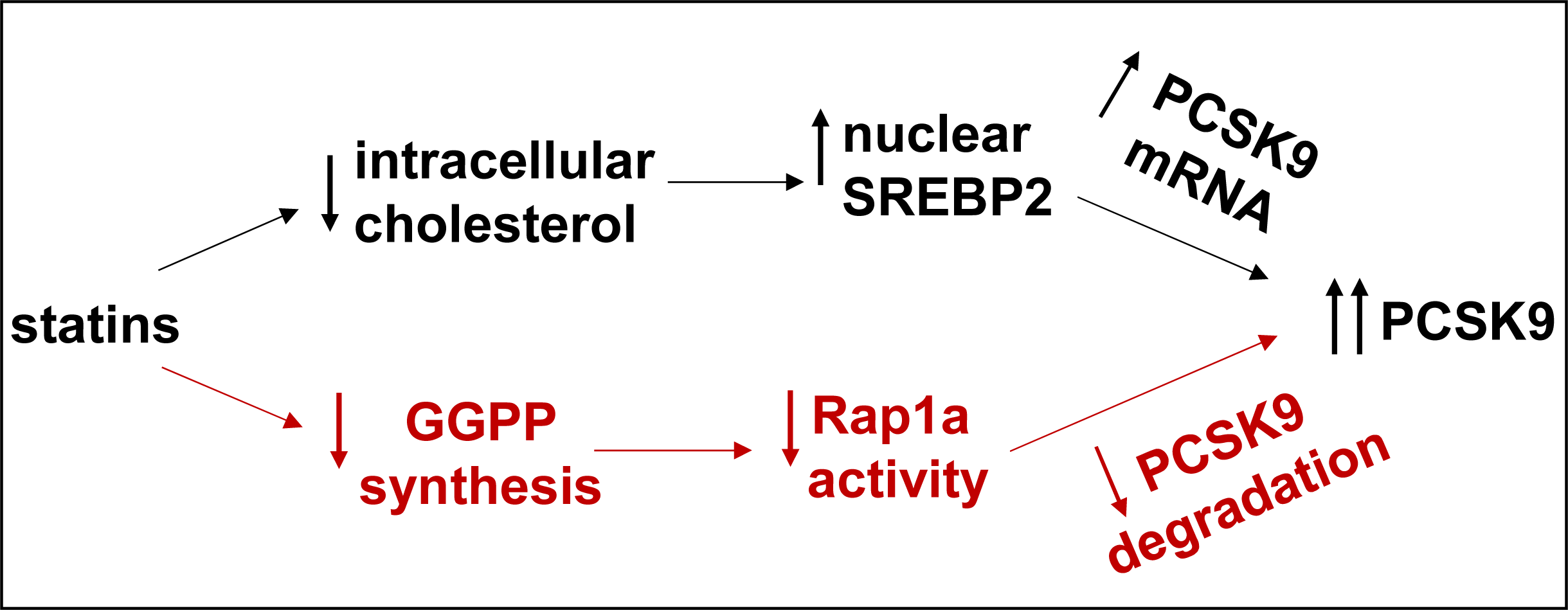
Proposed PCSK9 regulation by statins. Statins increase PCSK9 through increasing its mRNA via SREBP2 and inhibiting its degradation by decreasing the GGPP synthesis and Rap1a activity.

PCSK9 inhibitors are now used in the clinics and are well tolerated. Interestingly, genetic variants in *PCSK9* that mimic the effect of PCSK9 inhibitors are associated with increased risk for type 2 diabetes (T2D) development, raising the possibility that long term PCSK9 inhibition may result in new-onset T2D.^61–64^ Although this view has been challenged by many trials using anti-PCSK9 antibodies,^65,66^ a recent meta-analysis of randomized clinical trials of PCSK9 inhibitors reported a modest increase in plasma glucose and HbA1C with anti-PCSK9 antibodies.^67^ In this regard, it is tempting to speculate that activating Rap1a has the benefit of improving T2D as we recently reported that activating Rap1a improves glucose tolerance in mouse models of T2D.^36^

Our data suggest that Rap1a regulates PCSK9 via its downstream effectors, RhoA and ROCK and previous work revealed that treatment of mice with a ROCK inhibitor lowered plasma cholesterol.^68^ Interestingly, a recent study reported that statins increased RhoA activity in human lymphoblastoid cells, which inversely correlated with their cell surface LDLR protein.^54^ Moreover, the greater statin-induced RhoA levels were associated with reduced LDL-C lowering effect of statins in individuals from whom the cell lines were derived.^54^ In this regard, we postulate that our results provide a plausible explanation for the link between RhoA-ROCK and LDL-C metabolism and suggest that RhoA-ROCK mediated PCSK9 regulation may be the key underlying mechanism of this effect.

In summary, the work presented here further describes an essential role for Rap1a prenylation and activity in the regulation of plasma PCSK9 and LDL-C. Further, we provide evidence that GGPP supplementation and Rap1a activators can overcome the statin mediated PCSK9 induction and increase their efficacy. Our results may suggest the future use of combination therapy of Rap1a activators or GGPP add back with lower doses of statins, which would increase statins’ tolerability and decrease side effects without compromising efficacy.

## Acknowledgements

We thank Dr. Carol L. Williams (Medical College of Wisconsin) for Rap1a (wildtype) and the Rap1a-SAAX mutant plasmids and Dr. Ira Tabas and Dr. Xiaobo Wang (Columbia University) for providing liver tissues of AAV8-sh-RhoA or control AAV (Con)-treated mice fed with a high-cholesterol diet.

## Notes

**Funding sources:** This work was supported by National Natural Science Foundation of China (No.82300681) and Hunan Provencial Natural Science Foundation of China (No.2023JJ40856) to Y.W.; NIH grant (HL165129) to L.O.; and a Research Supplement to Promote Diversity in Science (23DIVSUP1071039) from American Heart Association to B.S.

### Competing Interest Statement

The authors have declared no competing interest.

## References

1. Abifadel M, Varret M, Rabes JP, Allard D, Ouguerram K, Devillers M, Cruaud C, Benjannet S, Wickham L, Erlich D, et al. Mutations in PCSK9 cause autosomal dominant hypercholesterolemia. Nat Genet. 2003;34:154–156. doi: 10.1038/ng1161

2. Horton JD, Cohen JC, Hobbs HH. Molecular biology of PCSK9: its role in LDL metabolism. Trends Biochem Sci. 2007;32:71–77. doi: 10.1016/j.tibs.2006.12.008

3. Shapiro MD, Tavori H, Fazio S. PCSK9: From Basic Science Discoveries to Clinical Trials. Circ Res. 2018;122:1420–1438. doi: 10.1161/CIRCRESAHA.118.311227

4. Sun H, Krauss RM, Chang JT, Teng BB. PCSK9 deficiency reduces atherosclerosis, apolipoprotein B secretion, and endothelial dysfunction. J Lipid Res. 2018;59:207–223. doi: 10.1194/jlr.M078360

5. Karagiannis AD, Liu M, Toth PP, Zhao S, Agrawal DK, Libby P, Chatzizisis YS. Pleiotropic Anti-atherosclerotic Effects of PCSK9 InhibitorsFrom Molecular Biology to Clinical Translation. Curr Atheroscler Rep. 2018;20:20. doi: 10.1007/s11883-018-0718-x

6. Arsenault BJ, Petrides F, Tabet F, Bao W, Hovingh GK, Boekholdt SM, Ramin-Mangata S, Meilhac O, DeMicco D, Rye KA, et al. Effect of atorvastatin, cholesterol ester transfer protein inhibition, and diabetes mellitus on circulating proprotein subtilisin kexin type 9 and lipoprotein(a) levels in patients at high cardiovascular risk. J Clin Lipidol. 2018;12:130–136. doi: 10.1016/j.jacl.2017.10.001

7. Momtazi-Borojeni AA, Sabouri-Rad S, Gotto AM, Pirro M, Banach M, Awan Z, Barreto GE, Sahebkar A. PCSK9 and inflammation: a review of experimental and clinical evidence. Eur Heart J Cardiovasc Pharmacother. 2019;5:237–245. doi: 10.1093/ehjcvp/pvz022

8. Bergmark B, O’Donoghue M, Murphy S, Kuder J, Ezhov MV, Ceska R, Gouni-Berthold I, Jensen HK, Tokgozoglu SL, Mach F, et al. PCSK9 INHIBITION AND AORTIC STENOSIS IN THE FOURIER TRIAL. Journal of the American College of Cardiology. 2020;75:2112–2112. doi: doi:10.1016/S0735-1097(20)32739-X

9. Weintraub WS, Gidding SS. PCSK9 Inhibitors: A Technology Worth Paying For? Pharmacoeconomics. 2016;34:217–220. doi: 10.1007/s40273-015-0355-y

10. Hardy J, Niman S, Pereira E, Lewis T, Reid J, Choksi R, Goldfaden RF. A Critical Review of the Efficacy and Safety of Inclisiran. Am J Cardiovasc Drugs. 2021. doi: 10.1007/s40256-021-00477-7

11. Goldstein JL, Brown MS. The LDL receptor. Arterioscler Thromb Vasc Biol. 2009;29:431–438. doi: 10.1161/atvbaha.108.179564

12. Baigent C, Keech A, Kearney PM, Blackwell L, Buck G, Pollicino C, Kirby A, Sourjina T, Peto R, Collins R, et al. Efficacy and safety of cholesterol-lowering treatment: prospective meta-analysis of data from 90,056 participants in 14 randomised trials of statins. Lancet. 2005;366:1267–1278. doi: 10.1016/S0140-6736(05)67394-1

13. Gitt AK, Drexel H, Feely J, Ferrières J, Gonzalez-Juanatey JR, Thomsen KK, Leiter LA, Lundman P, da Silva PM, Pedersen T, et al. Persistent lipid abnormalities in statin-treated patients and predictors of LDL-cholesterol goal achievement in clinical practice in Europe and Canada. Eur J Prev Cardiol. 2012;19:221–230. doi: 10.1177/1741826711400545

14. Wong ND, Chuang J, Zhao Y, Rosenblit PD. Residual dyslipidemia according to low-density lipoprotein cholesterol, non-high-density lipoprotein cholesterol, and apolipoprotein B among statin-treated US adults: National Health and Nutrition Examination Survey 2009-2010. J Clin Lipidol. 2015;9:525–532. doi: 10.1016/j.jacl.2015.05.003

15. Ridker PM, Mora S, Rose L. Percent reduction in LDL cholesterol following high-intensity statin therapy: potential implications for guidelines and for the prescription of emerging lipid-lowering agents. Eur Heart J. 2016;37:1373–1379. doi: 10.1093/eurheartj/ehw046

16. Leitersdorf E. Cholesterol absorption inhibition: filling an unmet need in lipid-lowering management. European Heart Journal Supplements. 2001;3:E17–E23. doi: 10.1016/s1520-765x(01)90108-7

17. Oni-Orisan A, Hoffmann TJ, Ranatunga D, Medina MW, Jorgenson E, Schaefer C, Krauss RM, Iribarren C, Risch N. Characterization of Statin Low-Density Lipoprotein Cholesterol Dose-Response Using Electronic Health Records in a Large Population-Based Cohort. Circ Genom Precis Med. 2018;11:e002043. doi: 10.1161/CIRCGEN.117.002043

18. Dubuc G, Chamberland A, Wassef H, Davignon J, Seidah NG, Bernier L, Prat A. Statins upregulate PCSK9, the gene encoding the proprotein convertase neural apoptosis-regulated convertase-1 implicated in familial hypercholesterolemia. Arterioscler Thromb Vasc Biol. 2004;24:1454–1459. doi: 10.1161/01.Atv.0000134621.14315.43

19. Awan Z, Seidah NG, MacFadyen JG, Benjannet S, Chasman DI, Ridker PM, Genest J. Rosuvastatin, proprotein convertase subtilisin/kexin type 9 concentrations, and LDL cholesterol response: the JUPITER trial. Clin Chem. 2012;58:183–189. doi: 10.1373/clinchem.2011.172932

20. Berthold HK, Seidah NG, Benjannet S, Gouni-Berthold I. Evidence from a randomized trial that simvastatin, but not ezetimibe, upregulates circulating PCSK9 levels. PLoS One. 2013;8:e60095. doi: 10.1371/journal.pone.0060095

21. Nozue T. Lipid Lowering Therapy and Circulating PCSK9 Concentration. J Atheroscler Thromb. 2017;24:895–907. doi: 10.5551/jat.RV17012

22. Welder G, Zineh I, Pacanowski MA, Troutt JS, Cao G, Konrad RJ. High-dose atorvastatin causes a rapid sustained increase in human serum PCSK9 and disrupts its correlation with LDL cholesterol. J Lipid Res. 2010;51:2714–2721. doi: 10.1194/jlr.M008144

23. Gallego-Colon E, Daum A, Yosefy C. Statins and PCSK9 inhibitors: A new lipid-lowering therapy. Eur J Pharmacol. 2020;878:173114. doi: 10.1016/j.ejphar.2020.173114

24. Miyoshi T, Nakamura K, Doi M, Ito H. Impact of Ezetimibe Alone or in Addition to a Statin on Plasma PCSK9 Concentrations in Patients with Type 2 Diabetes and Hypercholesterolemia: A Pilot Study. Am J Cardiovasc Drugs. 2015;15:213–219. doi: 10.1007/s40256-015-0119-2

25. Dong B, Wu M, Li H, Kraemer FB, Adeli K, Seidah NG, Park SW, Liu J. Strong induction of PCSK9 gene expression through HNF1alpha and SREBP2: mechanism for the resistance to LDL-cholesterol lowering effect of statins in dyslipidemic hamsters. J Lipid Res. 2010;51:1486–1495. doi: 10.1194/jlr.M003566

26. Jeong HJ, Lee HS, Kim KS, Kim YK, Yoon D, Park SW. Sterol-dependent regulation of proprotein convertase subtilisin/kexin type 9 expression by sterol-regulatory element binding protein-2. J Lipid Res. 2008;49:399–409. doi: 10.1194/jlr.M700443-JLR200

27. Hashemi M, Hoshyar R, Ande SR, Chen QM, Solomon C, Zuse A, Naderi M. Mevalonate Cascade and its Regulation in Cholesterol Metabolism in Different Tissues in Health and Disease. Curr Mol Pharmacol. 2017;10:13–26. doi: 10.2174/1874467209666160112123746

28. Miziorko HM. Enzymes of the mevalonate pathway of isoprenoid biosynthesis. Arch Biochem Biophys. 2011;505:131–143. doi: 10.1016/j.abb.2010.09.028

29. Wang M, Casey PJ. Protein prenylation: unique fats make their mark on biology. Nat Rev Mol Cell Biol. 2016;17:110–122. doi: 10.1038/nrm.2015.11

30. Gloerich M, Bos JL. Regulating Rap small G-proteins in time and space. Trends Cell Biol. 2011;21:615–623. doi: 10.1016/j.tcb.2011.07.001

31. Wittchen ES, Aghajanian A, Burridge K. Isoform-specific differences between Rap1A and Rap1B GTPases in the formation of endothelial cell junctions. Small GTPases. 2011;2:65–76. doi: 10.4161/sgtp.2.2.15735

32. Kitayama H, Sugimoto Y, Matsuzaki T, Ikawa Y, Noda M. A ras-related gene with transformation suppressor activity. Cell. 1989;56:77–84.

33. de Rooij J, Zwartkruis FJ, Verheijen MH, Cool RH, Nijman SM, Wittinghofer A, Bos JL. Epac is a Rap1 guanine-nucleotide-exchange factor directly activated by cyclic AMP. Nature. 1998;396:474–477. doi: 10.1038/24884

34. Bos JL, Rehmann H, Wittinghofer A. GEFs and GAPs: critical elements in the control of small G proteins. Cell. 2007;129:865–877. doi: 10.1016/j.cell.2007.05.018

35. Spolitu S, Okamoto H, Dai W, Zadroga JA, Wittchen ES, Gromada J, Ozcan L. Hepatic Glucagon Signaling Regulates PCSK9 and Low-Density Lipoprotein Cholesterol. Circ Res. 2019;124:38–51. doi: 10.1161/CIRCRESAHA.118.313648

36. Wang Y, Spolitu S, Zadroga JA, Sarecha AK, Ozcan L. Hepatocyte Rap1a contributes to obesity-and statin-associated hyperglycemia. Cell Rep. 2022;40:111259. doi: 10.1016/j.celrep.2022.111259

37. Wang X, Cai B, Yang X, Sonubi OO, Zheng Z, Ramakrishnan R, Shi H, Valenti L, Pajvani UB, Sandhu J, et al. Cholesterol Stabilizes TAZ in Hepatocytes to Promote Experimental Non-alcoholic Steatohepatitis. Cell Metab. 2020;31:969–986 e967. doi: 10.1016/j.cmet.2020.03.010

38. Lisowski L, Dane AP, Chu K, Zhang Y, Cunningham SC, Wilson EM, Nygaard S, Grompe M, Alexander IE, Kay MA. Selection and evaluation of clinically relevant AAV variants in a xenograft liver model. Nature. 2014;506:382–386. doi: 10.1038/nature12875

39. Wittchen ES, Worthylake RA, Kelly P, Casey PJ, Quilliam LA, Burridge K. Rap1 GTPase inhibits leukocyte transmigration by promoting endothelial barrier function. J Biol Chem. 2005;280:11675–11682. doi: 10.1074/jbc.M412595200

40. Wilson JM, Prokop JW, Lorimer E, Ntantie E, Williams CL. Differences in the Phosphorylation-Dependent Regulation of Prenylation of Rap1A and Rap1B. J Mol Biol. 2016;428:4929–4945. doi: 10.1016/j.jmb.2016.10.016

41. Basu D, Hu Y, Huggins LA, Mullick AE, Graham MJ, Wietecha T, Barnhart S, Mogul A, Pfeiffer K, Zirlik A, et al. Novel Reversible Model of Atherosclerosis and Regression Using Oligonucleotide Regulation of the LDL Receptor. Circ Res. 2018;122:560–567. doi: 10.1161/CIRCRESAHA.117.311361

42. Ozcan L, Wong CC, Li G, Xu T, Pajvani U, Park SK, Wronska A, Chen BX, Marks AR, Fukamizu A, et al. Calcium signaling through CaMKII regulates hepatic glucose production in fasting and obesity. Cell Metab. 2012;15:739–751. doi: 10.1016/j.cmet.2012.03.002

43. Gibbs BS, Zahn TJ, Mu Y, Sebolt-Leopold JS, Gibbs RA. Novel farnesol and geranylgeraniol analogues: A potential new class of anticancer agents directed against protein prenylation. J Med Chem. 1999;42:3800–3808. doi: 10.1021/jm9902786

44. Rashid S, Curtis DE, Garuti R, Anderson NN, Bashmakov Y, Ho YK, Hammer RE, Moon YA, Horton JD. Decreased plasma cholesterol and hypersensitivity to statins in mice lacking Pcsk9. Proc Natl Acad Sci U S A. 2005;102:5374–5379. doi: 10.1073/pnas.0501652102

45. Schuld NJ, Vervacke JS, Lorimer EL, Simon NC, Hauser AD, Barbieri JT, Distefano MD, Williams CL. The chaperone protein SmgGDS interacts with small GTPases entering the prenylation pathway by recognizing the last amino acid in the CAAX motif. J Biol Chem. 2014;289:6862–6876. doi: 10.1074/jbc.M113.527192

46. Palsuledesai CC, Distefano MD. Protein prenylation: enzymes, therapeutics, and biotechnology applications. ACS Chem Biol. 2015;10:51–62. doi: 10.1021/cb500791f

47. Xu N, Shen N, Wang X, Jiang S, Xue B, Li C. Protein prenylation and human diseases: a balance of protein farnesylation and geranylgeranylation. Sci China Life Sci. 2015;58:328–335. doi: 10.1007/s11427-015-4836-1

48. Gloerich M, Bos JL. Epac: defining a new mechanism for cAMP action. Annu Rev Pharmacol Toxicol. 2010;50:355–375. doi: 10.1146/annurev.pharmtox.010909.105714

49. Kim JG, Islam R, Cho JY, Jeong H, Cap KC, Park Y, Hossain AJ, Park JB. Regulation of RhoA GTPase and various transcription factors in the RhoA pathway. J Cell Physiol. 2018;233:6381–6392. doi: 10.1002/jcp.26487

50. Zhu Y, Casey PJ, Kumar AP, Pervaiz S. Deciphering the signaling networks underlying simvastatin-induced apoptosis in human cancer cells: evidence for non-canonical activation of RhoA and Rac1 GTPases. Cell Death Dis. 2013;4:e568. doi: 10.1038/cddis.2013.103

51. Li Z, Zhang J, Xue Y, He Y, Tang L, Ke M, Gong Y. Pitavastatin stimulates retinal angiogenesis via HMG-CoA reductase-independent activation of RhoA-mediated pathways and focal adhesion. Graefes Arch Clin Exp Ophthalmol. 2021;259:2707–2716. doi: 10.1007/s00417-021-05328-4

52. Akula MK, Ibrahim MX, Ivarsson EG, Khan OM, Kumar IT, Erlandsson M, Karlsson C, Xu X, Brisslert M, Brakebusch C, et al. Protein prenylation restrains innate immunity by inhibiting Rac1 effector interactions. Nat Commun. 2019;10:3975. doi: 10.1038/s41467-019-11606-x

53. Khan OM, Ibrahim MX, Jonsson IM, Karlsson C, Liu M, Sjogren AK, Olofsson FJ, Brisslert M, Andersson S, Ohlsson C, et al. Geranylgeranyltransferase type I (GGTase-I) deficiency hyperactivates macrophages and induces erosive arthritis in mice. J Clin Invest. 2011;121:628–639. doi: 10.1172/JCI43758

54. Medina MW, Theusch E, Naidoo D, Bauzon F, Stevens K, Mangravite LM, Kuang YL, Krauss RM. RHOA is a modulator of the cholesterol-lowering effects of statin. PLoS Genet. 2012;8:e1003058. doi: 10.1371/journal.pgen.1003058

55. Post A, Pannekoek WJ, Ponsioen B, Vliem MJ, Bos JL. Rap1 Spatially Controls ArhGAP29 To Inhibit Rho Signaling during Endothelial Barrier Regulation. Mol Cell Biol. 2015;35:2495–2502. doi: 10.1128/mcb.01453-14

56. Post A, Pannekoek WJ, Ross SH, Verlaan I, Brouwer PM, Bos JL. Rasip1 mediates Rap1 regulation of Rho in endothelial barrier function through ArhGAP29. Proc Natl Acad Sci U S A. 2013;110:11427–11432. doi: 10.1073/pnas.1306595110

57. Wang Y, Huang Y, Hobbs HH, Cohen JC. Molecular characterization of proprotein convertase subtilisin/kexin type 9-mediated degradation of the LDLR. J Lipid Res. 2012;53:1932–1943. doi: 10.1194/jlr.M028563

58. Nguyen MA, Kosenko T, Lagace TA. Internalized PCSK9 dissociates from recycling LDL receptors in PCSK9-resistant SV-589 fibroblasts. J Lipid Res. 2014;55:266–275. doi: 10.1194/jlr.M044156

59. Chen YF, Huang G, Wang YM, Cheng M, Zhu FF, Zhong JN, Gao YD. Exchange protein directly activated by cAMP (Epac) protects against airway inflammation and airway remodeling in asthmatic mice. Respir Res. 2019;20:285. doi: 10.1186/s12931-019-1260-2

60. Stokman G, Qin Y, Genieser HG, Schwede F, de Heer E, Bos JL, Bajema IM, van de Water B, Price LS. Epac-Rap signaling reduces cellular stress and ischemia-induced kidney failure. J Am Soc Nephrol. 2011;22:859–872. doi: 10.1681/ASN.2010040423

61. Lotta LA, Sharp SJ, Burgess S, Perry JRB, Stewart ID, Willems SM, Luan J, Ardanaz E, Arriola L, Balkau B, et al. Association Between Low-Density Lipoprotein Cholesterol-Lowering Genetic Variants and Risk of Type 2 Diabetes: A Meta-analysis. JAMA. 2016;316:1383–1391. doi: 10.1001/jama.2016.14568

62. Schmidt AF, Swerdlow DI, Holmes MV, Patel RS, Fairhurst-Hunter Z, Lyall DM, Hartwig FP, Horta BL, Hypponen E, Power C, et al. PCSK9 genetic variants and risk of type 2 diabetes: a mendelian randomisation study. Lancet Diabetes Endocrinol. 2017;5:97–105. doi: 10.1016/S2213-8587(16)30396-5

63. Ference BA, Robinson JG, Brook RD, Catapano AL, Chapman MJ, Neff DR, Voros S, Giugliano RP, Davey Smith G, Fazio S, et al. Variation in PCSK9 and HMGCR and Risk of Cardiovascular Disease and Diabetes. N Engl J Med. 2016;375:2144–2153. doi: 10.1056/NEJMoa1604304

64. Tcheoubi SER, Akpovi CD, Coppee F, Decleves AE, Laurent S, Agbangla C, Burtea C. Molecular and cellular biology of PCSK9: impact on glucose homeostasis. J Drug Target. 2022;30:948–960. doi: 10.1080/1061186X.2022.2092622

65. Colhoun HM, Ginsberg HN, Robinson JG, Leiter LA, Muller-Wieland D, Henry RR, Cariou B, Baccara-Dinet MT, Pordy R, Merlet L, et al. No effect of PCSK9 inhibitor alirocumab on the incidence of diabetes in a pooled analysis from 10 ODYSSEY Phase 3 studies. Eur Heart J. 2016;37:2981–2989. doi: 10.1093/eurheartj/ehw292

66. Blom DJ, Koren MJ, Roth E, Monsalvo ML, Djedjos CS, Nelson P, Elliott M, Wasserman SM, Ballantyne CM, Holman RR. Evaluation of the efficacy, safety and glycaemic effects of evolocumab (AMG 145) in hypercholesterolaemic patients stratified by glycaemic status and metabolic syndrome. Diabetes Obes Metab. 2017;19:98–107. doi: 10.1111/dom.12788

67. de Carvalho LSF, Campos AM, Sposito AC. Proprotein Convertase Subtilisin/Kexin Type 9 (PCSK9) Inhibitors and Incident Type 2 Diabetes: A Systematic Review and Meta-analysis With Over 96,000 Patient-Years. Diabetes Care. 2018;41:364–367. doi: 10.2337/dc17-1464

68. Abdali NT, Yaseen AH, Said E, Ibrahim TM. Rho kinase inhibitor fasudil mitigates high-cholesterol diet-induced hypercholesterolemia and vascular damage. Naunyn Schmiedebergs Arch Pharmacol. 2017;390:409–422. doi: 10.1007/s00210-017-1343-x

